# Modelling the probability of detecting mass mortality events

**DOI:** 10.1101/2021.01.16.426964

**Authors:** Jesse L Brunner, Justin M Calabrese

**Author notes:** **Corresponding Author**: Jesse L. Brunner, School of Biological Sciences, Washington State University, PO Box 644236, Pullman, Washington 99164.

## Abstract

While reports of mass mortality events (MMEs) are increasing in the literature, comparing the incidence of MMEs through time, among locations, or taxa is problematic without accounting for detection probabilities. MMEs involving small, cryptic species can be difficult to detect even during the event, and degradation and scavenging of carcasses can make the window for detection very short. As such, the number or occurrence rate of MMEs may often be severely underestimated, especially with infrequent observations. We develop a simple modeling framework to quantify the probability of detecting an MME as a function of the observation frequency relative to the rate at which MMEs become undetectable. This framework facilitates the design of surveillance programs and may be extended to correct estimates of the incidence of MMEs from actual surveillance data for more appropriate analyses of trends through time and among taxa.

## Introduction

Mass mortality events (MMEs), defined as “rapidly occurring catastrophic demographic events that punctuate background mortality levels” (Fey et al., 2015), can have profound impacts on populations, communities, and ecosystems. The growing number of reported MMEs suggests an increasing incidence of MMEs, which are often associated with natural and anthropogenic perturbations including disease, toxins, and multiple interacting stressors (Fey et al., 2015). While MMEs can produce spectacular levels of mortality, leaving hundreds to hundreds of thousands of carcasses in their wake, carcasses are scavenged, decompose, sink from view, or otherwise disappear (Santos et al., 2016), making MMEs difficult to detect without frequent surveillance (Santos et al., 2016; Kennedy et al., 2017). This is likely especially important for MMEs invovling soft-bodied species in environments with large numbers of scavengers or rapid rates of decay or flow through (Teixeira et al., 2013; Santos et al., 2016; Kennedy et al., 2017; Phelps et al., 2019). And MMEs involving small, rare, or cryptic species may be difficult to detect even at the peak of an event. There is thus a negative bias inherent to estimates of MMEs that is magnified in certain taxa and settings, making simple extrapolations from reported rates unreliable.

Unfortunately, quantifying detection probabilities and accounting for biases is difficult based on most published accounts. Basic details of how MMEs were detected, even whether they were discovered serendipitously or as part of routine surveillance, are usually absent (Fey et al., 2015, and references therein). The literature, however, does appear biased towards more easily detected species and accessible settings. For instance, half (52%) of the 460 reports used by Fey et al. (2015) involved fishes, perhaps because large numbers of carcasses floating near shore or in fishing areas are difficult to miss. Most of the 64 MMEs of mammals involved large-bodied species. All but five of the 27 MMEs in amphibians involved species associated with water bodies, where MMEs would be easier to detect, and the few reports in terrestrial environments involved intense surveillance efforts (Brem and Lips, 2008; Crawford et al., 2010; Attademo et al., 2011) or well-observed locations (Holland et al., 1990; Duff et al., 2011). Indeed, outside of systems frequently observed in research or other activities (e.g., fishing), most reports of MMEs seem to involve a large measure of coincidence.

While it is apparent that MMEs are detected imperfectly in many contexts and taxa, quantifying the degree of bias has so far been impossible. What is needed is a quantitative framework that identifies key determinants of MME detectability and makes clear their relationships. Such a framework could serve both as a basis for future statistical developments focused on estimating MME detection probability and as a means to standardize how MMEs are reported in the literature. Currently, no such framework exists. While there is an established literature on detection of carcasses (e.g., during road surveys), existing models focus on estimating steady state mortality or kill rates (e.g., Teixeira et al., 2013) and do not extend straightforwardly to detecting MMEs, which are sporadic, intense events. In this short communication, we therefore focus on building simple models to explore how MME detection varies with initial detectability, the frequency of surveillance, and change in detectability over time.

## Methods

We first make the simplifying assumption that a single mass mortality event occurs randomly, with uniform probability at any time, (0*, T*), during a season of length *T*. This avoids complications arising from potential dependence between events in terms of timing or detectability, but is realistic in situations where surveillance efforts or MMEs are strongly seasonal or involve such a large fraction of individuals that a subsequent event would be essentially undetectable. Second, we assume *n* observations are made regularly during the period starting at *t* = 0, then at intervals of *T/*(*n* − 1), and concluding at *t* = *T*.

We developed two versions of the model to reflect how detection probabilities might change through time. In the first model, the MME occurs instantaneously and has an initial (peak) detectability *α* ∈ [0, 1], but the probability of detection subsequently decays exponentially at rate *λ >* 0. Larger values of *α* represent MMEs that are easily detected by an observer, such as those involving large or abundant species in easily observed locations (e.g., charismatic megafauna in open fields or fishes near shore) whereas small values of *α* would correspond to MMEs of small, cryptic, or rare species in locations that are difficult to study (e.g., small shrews in dense forests or fossorial caecilians). Similarly, small, soft-bodied taxa that degrade quickly or that are rapidly scavenged or washed out of aquatic systems (e.g., Phelps et al., 2019) would have larger values of *λ*. While the probability of detecting animal carcasses, and thus MMEs, depends on myriad factors affecting carcass persistence and detectability (e.g., Korner-Nievergelt et al., 2015; Santos et al., 2016), exponential decay accounts for the general trend of declining detection probability as carcasses decay and are removed at a more or less constant rate (e.g., Regester and Whiles, 2006).

The second model assumes detection probability follows a Laplace (i.e., back-to-back exponential) distribution, such that detection probability first increases exponentially at rate *λ*, peaks at *α*, and then declines exponentially at rate *λ* thereafter. This model may be more representative of autocatalytic events such as epidemics of disease. Note that in both cases, we are using rescaled (to peak at *α*) probability functions as convenient functional forms that represent change in detection probability vs. time during the season, and are not assuming that detection probabilities are random variables.

For simplicity, we scale time in both models such that *T* = 1 and *λ* is defined relative to this period. In addition to eliminating the need to explore how *T* influences model behavior, this formulation also highlights the intuitive notion that it is not the absolute time between observations, but rather the time between observations, *T/*(*n* − 1), relative to the rate at which MMEs become undetectable, *λ*, that determines one’s chance of detecting an event. In any case, one can always rescale to an absolute rate (e.g., day^−1^) as *λ*_absolute_ = *λ*_scaled_*/T*, for *T >* 1.

### Exponential model

The probability of detecting a MME occurring at time *t* = *x* in the *n* equally spaced observations is one minus the product of the probabilities of *not* detecting the event at each observation. However, the event time *x* is a uniformly distributed random variable over the (scaled) season (0, 1), and to obtain the marginal probability of detection, which accounts for variation in event timing, we must integrate over the distribution of *x*. Therefore, the marginal probability of detecting a randomly occurring mortality event, assuming exponential decay of detection probability, is given by

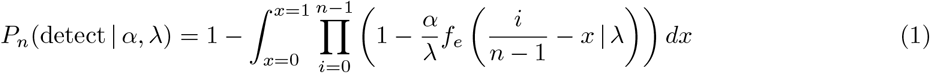

where

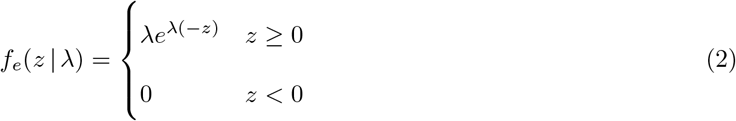

is the PDF of the exponential distribution.

Exact solutions to (1) can be constructed for particular values of *n*, for instance

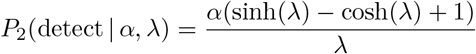

when *n* = 2 and

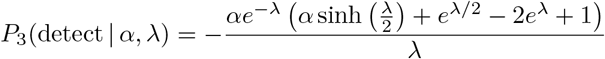

for *n* = 3, and so on. However, these exact expressions become cumbersome for larger *n*. Instead, the general solution (1) can be solved numerically for any *n* ≥ 2 (see Appendix A for example code).

### Laplace model

When detection probabilities follow a Laplace function, the probability of detecting a random event is:

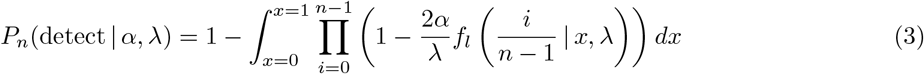

where

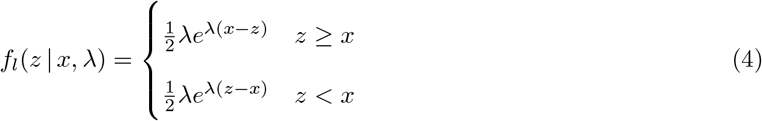

is the PDF of the Laplace distribution.

Exact solutions can again be constructed for particular values of *n*, such as

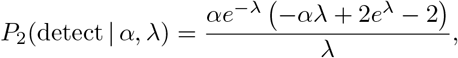

for *n* = 2 and

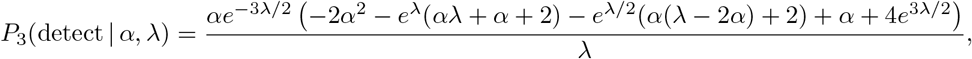

for *n* = 3. As for the exponential model, the exact expressions become unwieldy for larger *n*, but numerical solutions for (3) can be obtained (Appendix A).

## Results

For both models, events that are easily detected at their peak (high values of *α*) and that remain detectable over long periods (low values of *λ*) are more likely to be detected than those with low detectability at their peak (low *α*) or that quickly fade (high values of *λ*; Fig. 1). Similarly, events that become more detectable up to the peak of the event and then decline thereafter (Laplace model; Fig. 1) are, all else being equal, more easily detected than those that become instantly detectable and then decline (exponential model; Fig. 1). These results are intuitive, but it is striking how frequent observations must be to ensure a reasonable probability of detecting rapidly fading events (Fig. 2); five surveys during a season would guarantee a ≥ 80% chance of detecting a MME for only the two lowest decay rates (*λ* ≤ 2) we consider in Fig. 1 when the event is perfectly detectable at it’s peak (*α* = 1). If *α* = 0.5, at least twice the number of observations would be required to achieve the same level of confidence at a given level of *λ* (Fig. 2).

**Figure 1:**
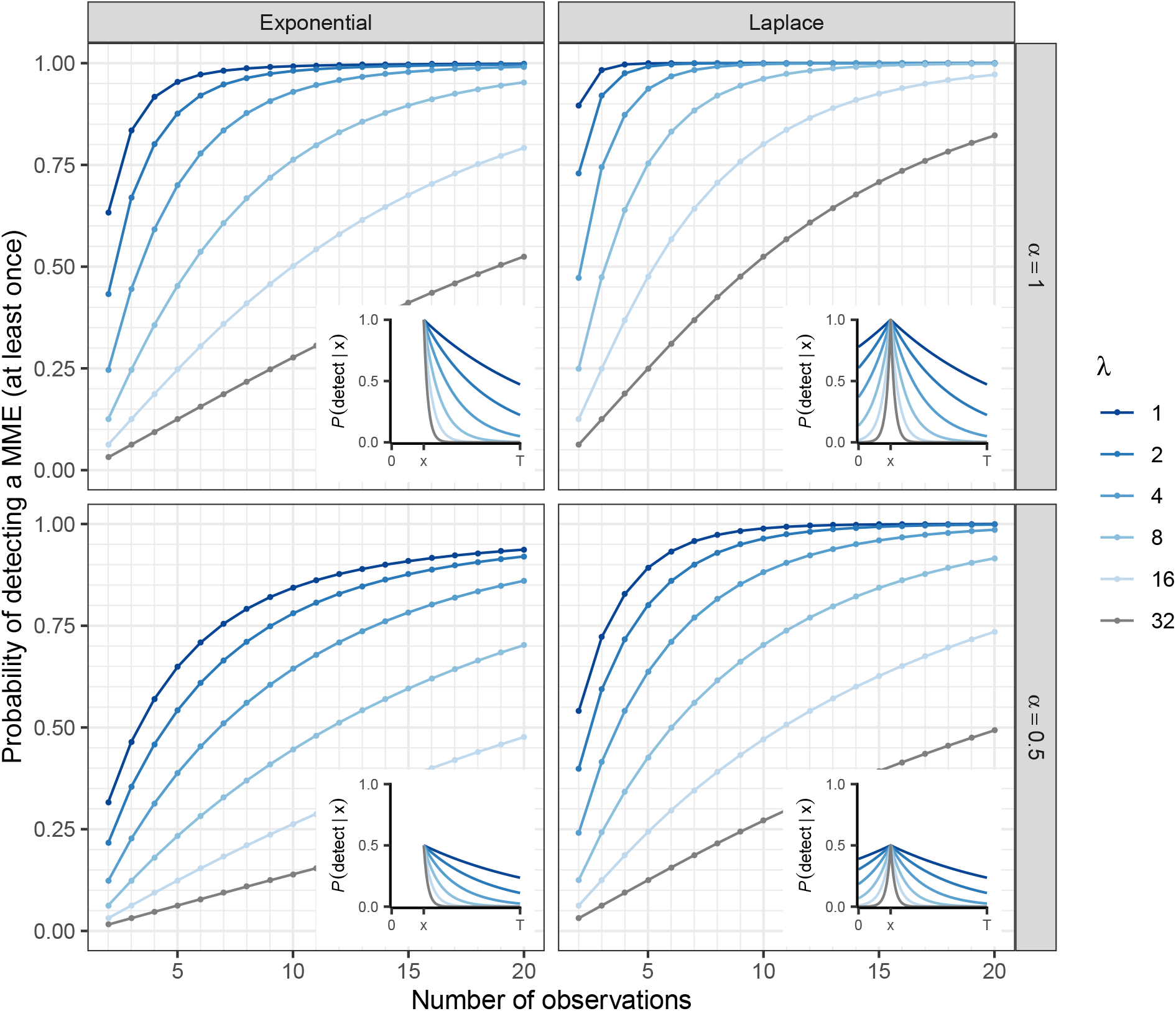
The marginal probability of detecting a mass mortality event (MME) occurring at some random time, *x*, as a function of the number observations made at regular intervals assuming either an exponential (left panel) or Laplace (right panel) form of detection probabilities over time. The inset graphs show how detection probabilities decline from a maximum of *α* = 1 (top row) and *α* = 0.5 (bottom row) at the peak of the MME at time *x*. Note that the rate of decline in detection probability, *λ*, is defined proportional to the length of the time in which MMEs can occur, *T*

**Figure 2:**
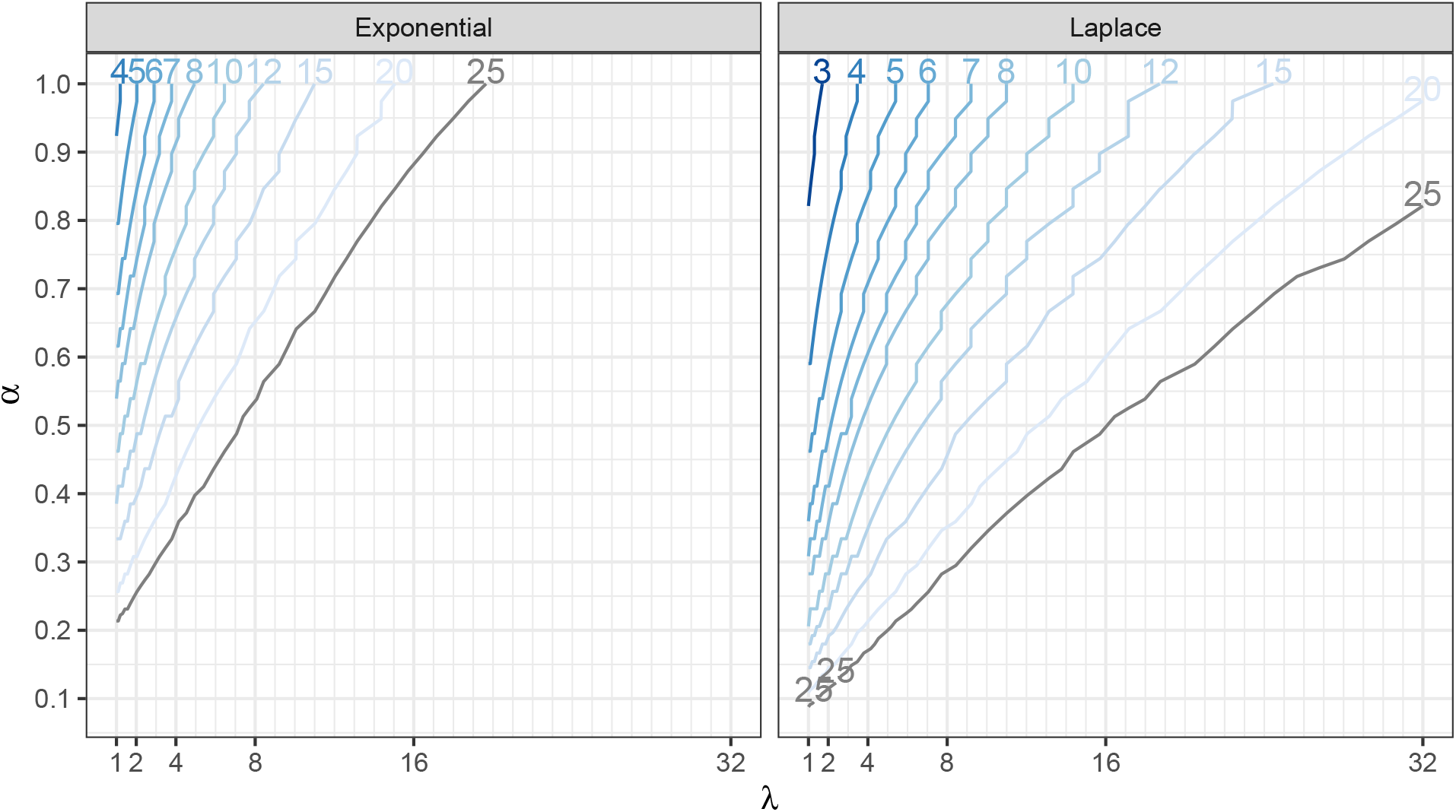
Isoclines of the number of regular observations (*n*; lines) required to achieve a ≥ 80% chance of observing a mass mortality event given particular values of the peak detection probability, *α*, and the rate at which detection probability declines from the event, *λ*, in the exponential (left panel) or Laplace (right panel) models.

As an example, consider the authors’ experience attempting to find *Ranavirus*-related MMEs in tiger salamanders inhabiting earthen cattle watering tanks in southern Arizona. They occur suddenly and sporadically, any time of year in this system, and can kill most or all aquatic stages. During one intense year of sampling 35 tanks were visited—seined or observed from shore—between three and 12 times (mean = 5.26, mode = 4), but just one MME was observed. Is this an accurate estimate? How likely were we to have detected MMEs given these frequencies of observations?

We used observations from prior years of 11 MMEs in eight tanks to estimate *λ* by fitting the exponential model of detection probability from the start of a MME (= first observation) to the binary detection data at each visit. This yielded *λ*_absolute_ ≈ 0.022 and thus *λ*_scaled_ = 0.022 × 365 ≈ 8. We also assumed *α* = 1 as these MMEs are large events and carcasses are often found floating or on shore. The exponential model with *λ* = 8 and *α* = 1 suggests our probability of detecting a MME in a tank varied from *P*_3_ = 0.25 with three visits to *P*_12_ = 0.83 with 12 visits, with an overall average of 0.44. Among the 16 tanks with four visits, the most common situation, there was a good chance that a MME would have gone undetected (1 − *P*_4_ = 0.64). While these are very rough estimates ignoring many details of the system (e.g., visits did not always occur at regular intervals, tanks varied in size and complexity), they suggest the actual frequency of MMEs was likely much greater than one in 35 tanks in the survey and that it is unsurprising the one MME we did detect was in a tank with more visits (*P*_8_ = 0.67). This rough analysis also suggests that to achieve an MME detection probability of 0.8, for example, would require *n* ≥ 11 observations (Fig. 2). If the peak probability of detection were even slightly lower, this number would grow (e.g., with *α* = 0.9, *n* ≥ 13).

## Discussion

Currently, there is no framework in the literature for quantifying MME detection probability, and this gap has limited progress in understanding how frequently MMEs occur in different systems. In this short communication, we have therefore developed a simple model that quantifies the probability of detecting MMEs as a function of how rapidly MMEs become undetectable, how often observations are made, and how detectable MMEs are at their peaks. This model provides a framework for both designing more effective surveillance programs and, potentially, standardizing empirical estimates of MME occurrence rates across systems and studies. It also highlights the key factors that empirical studies should attempt to report (i.e., *α, λ,* and *n*). At a minimum, reporting the sampling frequency (*n*) in the study should be encouraged, as well as relevant details of the focal area and search effort. It may also be possible to estimate detection probabilities at the peak of an event (*α*) and the change in detectability through time (*λ*) with, for instance, repeated surveys through time (e.g., Kennedy et al., 2017) or with replicate observers (e.g., Santos et al., 2016). Even coarse estimates, such as those we present for the *Ranavirus*-tiger salamander system, would allow researchers to identify important biases in observed rates of MMEs and design stronger surveillance programs.

The models developed here are intended to serve as a foundation upon which future developments can be built. For instance, while beyond the scope of this note, it would also be useful to expand our approach to accommodate covariates (e.g., search intensity, time of year), irregular observation intervals, and, potentially, multiple MMEs within a season, although the potential for dependence among sequential MMEs requires special attention. We also hope this work provides impetus for further empirical studies on how detection probabilities vary through time and among settings. Our initial work, however, suggests that the true rates of MMEs are likely substantially higher than what is observed, especially for small species in hot, difficult to study locations; that is, for much of the world’s biodiversity.

## Acknowledgments

JLB was supported by National Science Foundation grant 1754474.

## Appendix A

Here we present an R function to numerically solve equations (1) or (2) for any *n* ≥ 2. We first create an un-normalized Laplace probability density function with location parameter, mu, and the exponential growth and decay rate, lambda.

~~~
dlpl <- function(x, mu, lambda) exp(-abs(x-mu)*lambda)
~~~

Next, we create a function that returns the numerical solution from the probability of detecting an MME given either the exponential model (`mod=’Exp’`) or the Laplace model (`mod=’Lpl`). The parameter `lambda` must be greater than zero, `alpha` must be between zero and one, and `n` should be an integer ≥ 2. The function first creates an integrand function of the probability of not detecting an MME that is then integrated and subtrated from one to produce the probability of observing the MME at least once.

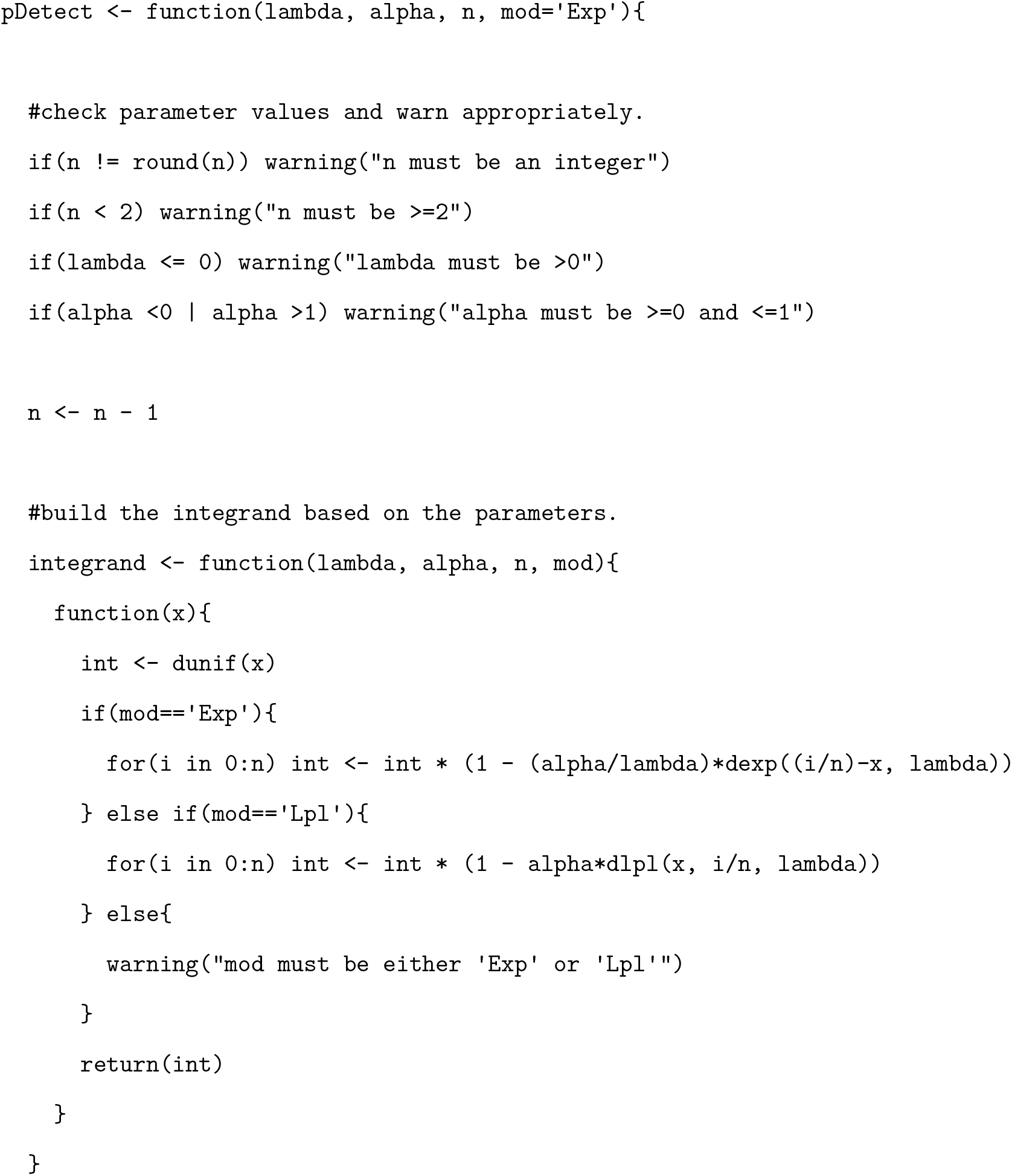
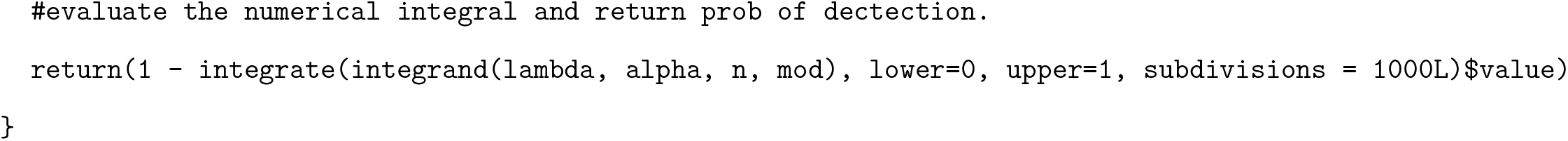

## Notes

### Competing Interest Statement

The authors have declared no competing interest.

